# Subcellular visualization of the distribution of atmospheric dinitrogen fixed by *Gluconacetobacter diazotrophicus* bacteria in maize

**DOI:** 10.1101/2024.04.29.591714

**Authors:** G. McMahon, S. Rey, K. Moore, G. Greenidge, D. Patel, E.H. Murchie, D. Dent, E. Cocking

**Affiliations:** National Physical Laboratory, National Centre of Excellence for Mass Spectrometry Imaging; Teddington, TW11 0LW, United Kingdom; National Physical Laboratory, Biometrology Group; Teddington, TW11 0LW, UK; University of Manchester, Photon Science Institute and Department of Materials; Manchester, M13 9PL, United Kingdom; University of Nottingham, School of Biosciences; Nottingham, LE12 5RD, United Kingdom; Azotic Technologies Ltd.; Dunnington, YO19 5SN, United Kingdom; ADAS Gleadthorpe; Mansfield, NG20 9PD, United Kingdom; Sustainable Nitrogen Foundation; Thetford, IP25 7HB, United Kingdom

**Author notes:** **Corresponding Author:** Dr. Greg McMahon, **Email:**. **Author Contributions:** Conceptualization: EC, DD, GM; Methodology: EC, RF, PH, PS, DP, MS; Investigation: RF, PH, PS, DP, MS, GM, SR, GG; Visualization: GM, GG, EM; Funding acquisition: EC, DD, DG; Writing – original draft: GM, EC, DD; Writing – review & editing: GM, KM, GG, DD, EM, EC. **Competing Interest Statement:** D. Dent and D. Patel were employees of Azotic Technologies Ltd. at the time some of the presented work was performed. D. Dent and the family of E. Cocking continue to own shares in Azotic Technologies Ltd.

**Keywords:** nitrogen fixation, maize, Gluconacetobacter diazotrophicus, NanoSIMS, chloroplasts

## Abstract

Plants normally obtain the nitrogen required for growth through their roots, often after application of synthetic fertilizer to the soil, at great cost to the environment and climate. Inoculation of plant seeds with nitrogen-fixing bacteria is a promising alternative means of supplying plants the nitrogen they require in an environmentally friendly manner. When maize seeds inoculated with nitrogen-fixing *Gluconacetobacter diazotrophicus (Gd)* are grown for two weeks in a ^15^N_2_ air environment, nanoscale secondary ion mass spectrometry (NanoSIMS) imaging shows the distribution of fixed nitrogen with subcellular resolution, with the majority being incorporated heterogeneously into chloroplasts. Chloroplasts, as the chief energy source that drives plant growth via photosynthesis, are vital for healthy plant growth and these results help explain the observations of enhanced growth rates in plants containing this nitrogen fixing bacteria. The methodology provides a template upon which more powerful, correlative studies combining genomic and/or spatial transcriptomic methods may be based.

## Introduction

Biological nitrogen fixation is exclusively performed by a few species of prokaryotes (bacteria and archaea) that possess the enzyme nitrogenase (*1*) which enables the reduction of N_2_ to ammonia. Plants and other eukaryotes do not possess nitrogenase and are therefore unable to fix nitrogen. They require fertilization with fixed nitrogen for their optimum growth and yield. However, the extensive global use of synthetic nitrogen fertilizers has not only resulted in the pollution of aquatic systems by soluble nitrates, with associated eutrophication and health hazards, but through soil bacterial activity, the ammonia is converted to nitrous oxide - a potent greenhouse gas associated with photochemical smog, fine particulate pollution, ecosystem acidification and climate change (*2*).

Legume crops, however, are able to establish colonies of symbiotic intracellular nitrogen-fixing rhizobia bacteria in their root nodules (*3*). In addition to legume crops, nitrogen fixing *Gluconacetobacter diazotrophicus* (*Gd*) bacteria were discovered in 1988 in the roots of Brazilian sugarcane plants (*4*). *Gd* is a highly versatile diazotroph, having one of the largest clusters of nitrogen fixing genes found in any diazotroph (*5*). It is able to fix nitrogen over a broad range of oxygen concentrations as well as protect the sensitive nitrogenase enzyme from oxygen inhibition through the interplay of a number factors relating to sucrose, colony structure, the levan extrapolysaccarharide, the detoxification of reactive oxygen species as well as control of oxygen through its respiratory pathway (*6-8*) to enable amino acid, chlorophyll and protein synthesis. Inoculation of plants with *Gd* bacteria and other intracellular nitrogen fixing bacteria is rapidly becoming a priority with potential to substitute for environmentally damaging synthetic nitrogen-based fertilizers. However, much remains to be learned, for example the exact mechanism by which bacteria enter the roots of plants after inoculation of seeds and how they contribute to growth. The aerobic endophytic diazotroph *Gd* (5541) isolated from sugarcane has been shown to possess enzymes such as endoglucanase, endopolymethylgalacturonase and endoxyloglucanase that enable bacterial penetration of plant cell walls (*9*). This behavior is somewhat similar to the interaction of rhizobia in root nodule cells (*10*), but without any nodule formation.

The techniques used to demonstrate nitrogen fixation in rhizobia have all been used to demonstrate nitrogen fixation by *Gd* in colonized plants, and evidence that nitrogen has been transferred from the atmosphere to the plant using stable isotope labelled ^15^N_2_ experiments combined with bulk isotope ratio mass spectrometer measurements (*5, 8, 11*) have been reported. However, these experiments lack any spatial data regarding ^15^N incorporation. In this study, we use Nanoscale secondary ion mass spectrometry (NanoSIMS) imaging to directly visualize the distribution of fixed nitrogen in various cellular organelles such as chloroplasts with subcellular resolution (down to ∼50 nm for instrumental conditions used).

## Results

Isotope ratio analysis by NanoSIMS of a control sample yielded a nitrogen isotope ratio of .0039 (SD = .000420, n = 40). This value is indeed (weakly) statistically significantly different from the theoretical natural ratio value taken as .0037 at the 95% confidence level (p = .0448; Wilcoxan signed rank test). However, the results that follow for the experimental samples show exceptionally high enrichments in ^15^N such that the small deviation of the control sample from the theoretical natural value will not impair interpretation of the data. Example raw NanoSIMS images for ^12^C^14^N, ^31^P, ^32^S and the derived ^12^C^15^N/^12^C^14^N image for a leaf taken from a plant grown in the ^15^N_2_ atmosphere are shown in Fig. 1. Significant uptake of ^15^N into chloroplasts (shown by the white arrows in Fig. 1a) are clearly observed in the nitrogen isotope ratio images in Fig. 1e and 1f. The cell nucleus, shown by the red arrow in Fig. 1a and identified through the high ^31^P signal (Fig. 1b) contributed by the DNA, has also received ^15^N, but to a lesser extent compared to the chloroplasts. This type of enrichment was consistent across the whole leaf (Fig. 1h). The data showed no obvious trend in the nitrogen isotope ratio value measured in the chloroplasts (n = 322) across the length of the leaf except for the sample taken 1 cm from the tip of the leaf, where the highest ratio values were consistently measured across several analytical sessions conducted over a time span of several months.

**Figure 1.**
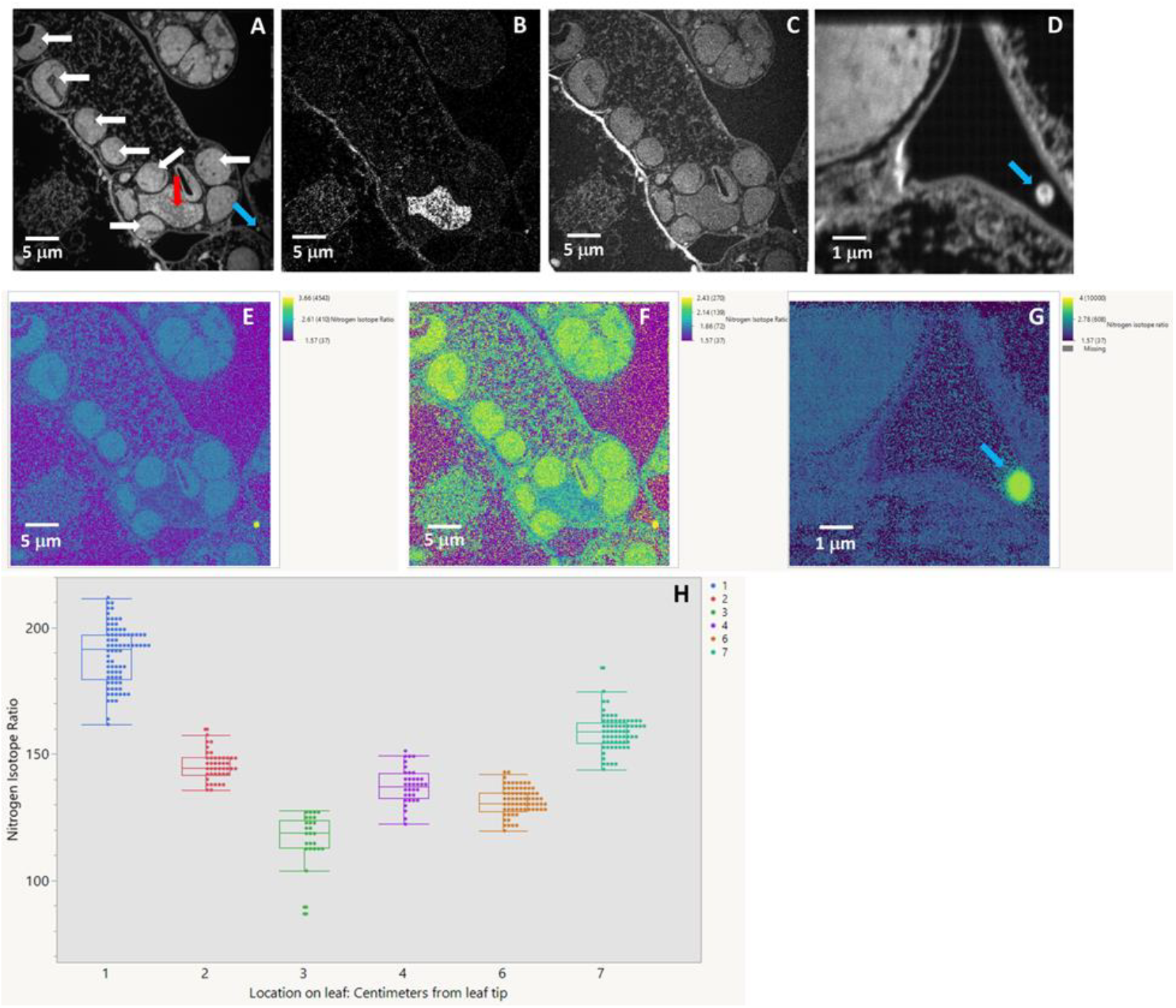
Raw NanoSIMS images from maize leaf for (a) ^12^C^14^N, (b) ^31^P, (c) ^32^S, (d) ^12^C^14^N, (e) the ^12^C^15^N/^12^C^14^N ratio image (log scale, full data range), (f) ^12^C^15^N/^12^C^14^N ratio image (log scale, reduced ratio scale maximum), (g) ^12^C^15^N/^12^C^14^N ratio image (log scale, full data range) and (h) distribution of chloroplast nitrogen isotope ratio values across leaf. Actual ratio values are shown in brackets in the log spectral scale. White arrows point to chloroplasts observed in the image. Red arrow points to cell nucleus. Blue arrow points to example of single bacteria-like structure with exceptionally high nitrogen isotope ratio.

The association between the nitrogen isotope ratio in the chloroplasts, nuclei and xylem cell walls across the entire leaf was also studied, irrespective of position between tip and stem. Regions of interest (ROIs) from the chloroplasts (n = 322), cell nuclei (n = 17), and xylem cell walls (n = 86) were generated and the nitrogen ratios from each region measured. The chloroplasts had significantly higher (p < .0001) nitrogen isotope ratio values compared to those measured in the nuclei and xylem cell walls, the latter of which were only slightly higher than the natural value.

An additional interesting observation was made within individual chloroplasts when analyzed at higher spatial resolution as shown in Fig. 2. The structure within the chloroplast is clearly visible within the ^12^C^14^N image (Fig. 2a), with the darker regions (arrowed) showing a granal lamellae type structure indicative of thylakoid membranes. These regions, which contain components of light harvesting and electron transport, had significantly less nitrogen (p < .0001, Fig. 2c) than the stromal regions where the more nitrogen demanding soluble Calvin-Benson cycle enzymes are located, including Rubisco (*13*), clearly observed in the nitrogen isotope ratio HSI image shown in Fig. 2b.

**Figure 2:**
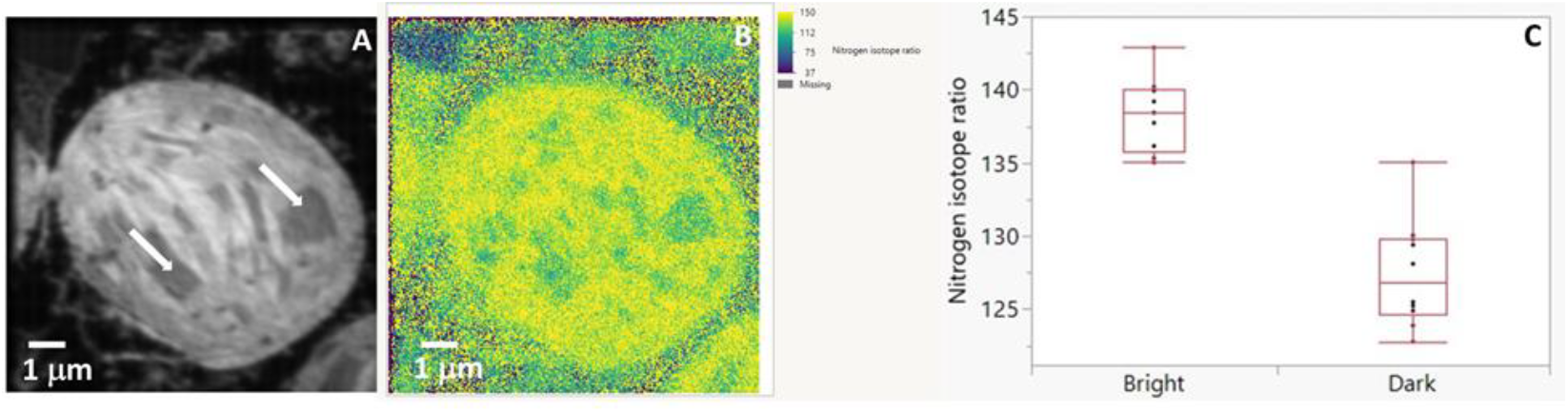
(a) High spatial resolution ^12^C^14^N image of a single chloroplast showing internal structures typical of thylakoid membranes (white arrows). (b) The nitrogen isotope ratio image demonstrates that these regions have assimilated significantly less fixed nitrogen (p < .0001; Fig. 3c) than the surrounding stroma where Rubisco is located.

We also sporadically observed areas with spectacular enrichment in ^15^N not associated with chloroplasts but with *Gd*-sized structures believed to be individual bacteria. A typical example is shown in Fig. 1d and 1g. These regions show immense enrichment in ^15^N (∼10,000% to over ∼27,000% increase above natural ratio). Note they are adjacent to areas with isotope ratio values barely above natural, acting as an internal control and indicating that these extremely high values are not an artefact of instrumental setup or detector miscalibration. Curiously, these regions also showed high sulphur counts in the raw image data (Fig. 1c), which could be explained by the presence and activity of the sulphur-rich nitrogenase enzyme, a necessary ingredient for nitrogen fixation. This may be a function of the symbiotic relationship with the plant, where the energy produced by photosynthesis is transferred to the bacteria to drive the nitrogen fixation reaction.

## Discussion

The methodology and data presented in this paper provide a means for directly observing the transport of *Gd* fixed nitrogen at subcellular resolution in maize plant leaves, driving biosynthesis and growth using nitrogen sourced from air, rather than synthetic fertilizer. Though NanoSIMS imaging has been used previously to study transport of gaseous dinitrogen fixed by cyanobacteria located within cyanobacteria cells in feather mosses in a boreal forest, no indication was given that it was being taken up by the chloroplasts (*14*), while Tarquinio *et al* (*15*) did observe high uptake in chloroplasts in seagrass, but in their case the administration of ^15^N was through a ^15^N-labeled amino acid solution rather than gaseous dinitrogen.

The findings reported in the present study may provide the basis of detailed agronomic analysis of the extent of fixed nitrogen fertilizer needs and yield responses in autonomous nitrogen-fixing maize plants and may also form part of more powerful correlative studies that also could include genomic and/or spatial transcriptomic analyses.

## Materials and Methods

Details regarding the materials and methods for the inoculation of maize seeds with *Gd* bacteria and growth of seedlings for 2 weeks in a ^15^N_2_ atmosphere are provided in the supplementary materials. Leaf sections were prepared for NanoSIMS by chemical fixation of hand sectioned pieces of the first leaf in 2% (v/v) glutaraldehyde in 0.1M sodium phosphate buffer, pH 7.0, for 24h at 48°C, dehydrated with graded ethanol, and embedded in LR White medium grade acrylic resin (Agar, UK) and sectioned at 500 nm thickness with a Leica Ultramicrotome and a Diatome Histo diamond knife. Sections for NanoSIMS were retrieved from the knife boat and deposited on a 5mm x 5mm silicon wafer.

NanoSIMS imaging was performed using a Cameca NanoSIMS 50L (Gennevilliers, France) using a Cs^+^ primary ion beam (16 keV impact energy/ion) with simultaneous detection of negative ^12^C^-, 16^O^-, 12^C^14^N^-, 12^C^15^N^-, 31^P^-^ and ^32^S^-^. The incorporation of ^15^N label is observed in the derived ^12^C^15^N/^12^C^14^N isotope ratio images. Further details pertaining to instrumental parameters are given in the supplementary data and data analysis methods have been previously described (12).

## Acknowledgments

This work is dedicated to the memory of the truly inspirational and gifted scientist, Prof. Ted Cocking. GM acknowledges Dr. G. Trindade and Dr. N. Belsey for their critical internal NPL reviews of the paper. The authors would also like to acknowledge M. Swaine, P. Stone and D. Green for technical assistance.

The funding for this work was provided by the NPL Measurement for Recovery program, funded by the former UK Department of Business, Energy and Industrial Strategy.

## Data Availability

All data available at https://doi.org/10.5281/zenodo.8315024

## Supplemental File

### Materials and Methods

Seeds of maize (Forage maize, Kaspian variety, Hunt Seeds) were surface sterilized in suitably diluted ‘Domestos’ bleach, c. 5% sodium hypochlorite (Lever Faberge’, Kingston-upon-Thames, UK), and rinsed in sterile water. Surface sterilized maize seeds were germinated on 15 ml of 0.8% (w/v) water agar in 9 cm Petri dishes (10 seeds/dish) for 3–4 d in the dark at 28°C and inoculated after transfer to the Murashige and Skoog (MS) agar medium.

For inoculation, an aqueous suspension of the *Gd* was prepared to give an optical density at 600 nm of 1.1, or ∼ 109 colony forming units (CFU) per milliliter. The number of CFU was determined by serial dilution, plating on ATGUS medium (with antibiotics as appropriate) and counting bacterial colonies after 4 d incubation in Petri dishes (28°C, dark). Seedlings growing in jars on MS agar medium were inoculated with 1 ml aliquots of the *Gd* around the bases of the stems with the exception of the control samples, where only distilled water was used. At germination, the seedlings were transferred to long test tubes with media having high sucrose but no nitrogen nutrient. Natural air was removed with vacuum from the test-tubes and replaced with a 70:30 mixture of O_2_ and ^15^N_2_ to mimic the air environment. Thus, only ^15^N_2_ was present for *Gd* to fix. The seedlings were left to grow for two weeks.

The NanoSIMS instrument used in this study (NanoSIMS 50L, Cameca, Gennevilliers, France) generates quantitative mass images at high spatial (down to ∼35-50 nm) and mass resolution (m/Dm ∼10,000) from which nitrogen isotope ratio images can be derived by simultaneously acquiring images from the light and heavy isotopes of negative secondary cyanide ions (^12^C^14^N^-^ and ^12^C^15^N^-^) produced by the bombardment of the samples with a primary cesium ion beam and simply dividing the two images. Isotope ratio values can be extracted from the images using a region of interest analysis, thus enabling the quantitative evaluation of fixed nitrogen incorporation into specific plant organelles. We raster scanned a 16 keV impact energy Cs^+^ primary ion beam across the sample and collected negatively charged secondary ions. The primary ion beam current ranged from ∼ 2 pA for the largest field sizes (50 mm x 50 mm) to < 0.5 pA for the smallest field sizes (5 mm x 5 mm). The beam current was largely dictated by the selected size of the D1 aperture (D1-2 = 300 mm diameter yielding ∼ 2 pA primary beam current; D1-5 = 100 mm diameter yielding < 0.5 pA primary beam current) and primary lens 1 (L1) voltage. For the highest spatial resolution images, L1 was set at 7500V and D1-5 selected. For all other images, L1 was set at 2000V. Dwell time per pixel ranged from 2-5 ms/pixel. After checking the pulse height distributions on all detectors, they were then positioned along the magnet radius to measure ^12^C, ^16^O, ^12^C_2_, ^12^C^14^N, ^12^C^15^N, ^31^P and ^32^S. Mass resolving power (MRP) was routinely > 7000 m/Dm, achieved by use of an entrance slit 30 mm in width by 180 mm in height (ES3) and an exit slit 50 mm in width by 1600 mm in height Data analysis was performed using the OpenMIMS plugin for ImageJ as described in (*12)*.

Statistical analysis was performed using JMP 17 Pro software. Normality of data was evaluated using Shapiro-Wilk test. Results of these tests indicated non-parametric testing (ie. data deviated from a normal distribution) would be the most appropriate. Thus, to compare the medians of the nitrogen isotope ratios of the various organelles, a Steel-Dwass test was used.

